# An assessment of Causal Inference in visual-inertial traveled distance estimation

**DOI:** 10.1101/357087

**Authors:** K.N. de Winkel, D. Diers, M. Laächele, H.H. Buülthoff

## Abstract

Recent work indicates that the central nervous system assesses the causality of visual and inertial information in the estimation of qualitative characteristics of self-motion and spatial orientation, and forms multisensory perceptions in accordance with the outcome of these assessments. Here, we extend the assessment of this Causal Inference (CI) strategy to the quantitative domain of traveled distance. We present a formal model of how stimuli result in sensory estimates, how percepts are constructed from sensory estimates, and how responses result from percepts. Starting with this formalization, we derived probabilistic formulations of CI and competing models for perception of traveled distance.

In an experiment, participants (n=9) were seated in the Max Planck Cablerobot Simulator, and shown a photo-realistic virtual rendering of the simulator hall via a Head-Mounted Display. Using this setup, the participants were presented with various unisensory and (incongruent) multisensory visual-inertial horizontal linear surge motions, differing only in amplitude (i.e., traveled distance). Participants performed both a Magnitude Estimation and a Two-Interval Forced Choice task.

Overall, model comparisons favor the CI model, but individual analysis shows a Cue Capture strategy is preferred in most individual cases. Parameter estimates indicate that visual and inertial sensory estimates follow a Stevens’ power law with positive exponent, and that noise increases with physical distance in accordance with a Weber’s law. Responses were found to be biased towards the mean stimulus distance, consistent with an interaction between percepts and prior knowledge in the formulation of responses. Magnitude estimate data further showed a regression to the mean effect.

The experimental data did not provide unambiguous support for the CI model. However, model derivations and fit results demonstrate it can reproduce empirical findings, arguing in favor of the CI model. Moreover, the methods outlined in the present study demonstrate how different sources of distortion in responses may be disentangled by combining psychophysical tasks.

## 1 Introduction

Humans, and many other animals alike, are able to form estimates of displacement of self-motion by integration of sensory information on translations and rotations over time; an ability that has become known as path integration [1, 2]. In humans, acceleration information from the body’s inertial sensors provides su cient information to perform this task [3, 4, 5, 6], and experiments performed in virtual reality [7, 8] have shown that humans can also form representations of displacement on the basis of visual optic flow information in isolation from inertial information. Consequently, the visual and inertial sensory modalities appear to provide the Central Nervous System (CNS) with redundant information on displacement. However, our conscious experience of it appears to be singular. It thus appears that the CNS applies some rule to distill a single supramodal internal representation out of multisensory information.

The mechanisms by which multisensory information on displacement may be processed have been the subject of a number of previous studies. Exploratory studies have found that multisensory estimates of displacement were somewhere in between the displacements of the unisensory cues; suggestive of multisensory interactions [9, 10, 11]. The studies’ findings however differed with respect to the relative weights attributed to each sensory modality: [9] found that inertial information was attributed a larger weight than visual information, whereas [10, 11] found the opposite. To reconcile these findings with contemporary literature on multisensory integration (e.g., [12, 13]), it was proposed that displacement estimation may be achieved by the CNS in a process that resembles Bayesian or Maximum-Likelihood Estimation[14]. According to these models, multisensory information is weighted according to its reliability and subsequently summed; differences between studies in the observed weightings of sensory estimates would ultimately be the result of differences in the reliability of the stimuli. In later work, sensory weightings were indeed found to be consistent with predictions of such modeling [15].

However, an axiomatic multisensory integration is problematic when applied to motion perception because visual information is inherently ambiguous with respect to its causality: identical patterns of visual stimulation result from self-motion through an environment as from the opposite motion of the environment relative to the observer (object motion). It is therefore plausible that the CNS assesses the causality of the visual and inertial information in the process of forming perceptions of self-motion.

The observations noted above translate to models of multisensory perception: the CNS could choose estimates provided by a single sensory system as the dominant source of information (Cue Capture, CC[16]; it could alternate between them on a trial-by-trial basis (Switching Strategy, SS[17]); it could combine estimates from different sensory systems into a single, integrated estimate by taking a weighted sum (Forced Fusion, FF[12, 13]); or it could favor different strategies, depending on an assessment of the causality of the visual and inertial information. Such an intermediate strategy that has been the focus of several recent studies on perception is Causal Inference (CI)[18, 19].

In previous work, we obtained evidence that the CNS performs CI in the estimation of heading of visual-inertial horizontal linear translations[20, 21], and in the estimation of verticality[22]. Here, we extend the assessment of the applicability of the CI model from the qualitative domain of heading and orientation[23] to the quantitative domain of traveled distance. Participants were seated in the Max-Planck CableRobot simulator, and wore an HMD that showed them a photo-realistic stereoscopic virtual version of their actual surroundings. Using this setup, we presented participants with combinations of visual and inertial horizontal translational (surge) motions, with various inter-sensory discrepancies in stimulus magnitude (i.e., traveled distance). Participants performed both a Magnitude Estimation (ME) and Two-Interval Forced Choice (2IFC) task. We fitted the CC, SS, FF, and CI models and compared them on the basis of their ability to explain the experimental data. Moreover, by fitting the models to the data of both tasks simultaneously, we disentangled sources of distortion in the responses; allowing us to quantify sensory biases as well as top-down e ects of prior knowledge.

## 2 Methods and Materials

### 2.1 Ethics Statement

The experimental protocol was approved by the ethical commission of the medical faculty of the Eberhard-Karls University in Tübingen, Germany, reference number 603/2017BO2.

### 2.2 Participants

In total, 17 people took part in the experiment. Out of these people, four were unable to perform the task due to motion sickness that occurred shortly after the start of the experiment. For four others only ME data was obtained, which was insu cient data for the present analysis. For the remaining nine participants, a complete dataset was obtained. Of these participants, three were female; six were male. Their average age was 28.33 years, with a standard deviation of 4.74 years. Participants were compensated for their time at a rate of €8 per hour.

### 2.3 Setup

Visual stimuli were presented using a Vive HMD (HTC, New Taipei City, Taiwan) displaying 1080 × 1200 pixels per eye at 90fps. The HMD was used to show participants ‘Alternative Reality’ (AR) motion stimuli, which consist of photo-realistic stereoscopic renderings of linear motion trajectories through a virtual model of participants’ actual surroundings (i.e., the simulator hall). The virtual hall model was created from a laser scan of the actual hall using photogrammetry. The entire scene was scaled according to actual dimensions. A view of the virtual simulator hall is shown in the left panel of Figure 1. Head motions were tracked and reproduced in the virtual environment using an OptiTrack V120 Trio optical tracking system (NaturalPoint Inc., Corvallis, OR, USA) that was mounted to the structure of the simulator cabin. The right panel in Figure 1 shows a (monoscopic) view from the simulator cabin, as seen by a participant.

**Fig. 1:**
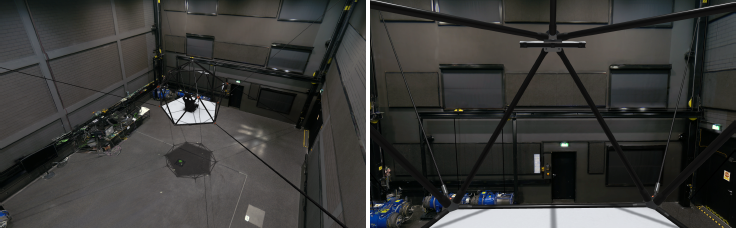
Left panel: view of the virtual simulator hall from above. Right panel: a monoscopic version of a participant’s view during the experiment. The image shows the view of the virtual hall from the cabin.

To generate inertial stimuli, we used the MPI CableRobot Simulator[24]. The CableRobot simulator features an icosahedron-shaped cabin that can be moved around within a workspace of 5 × 7 × 4m. Motion of the cabin is achieved using an array of eight motors that pull eight respective cables. Each cable is connected to the cabin via several pulleys placed throughout the room and winches that are directly actuated by the motors.

The simulator cabin is equipped with an automotive style bucket seat (RECARO, GmbH, Stuttgart, Germany). Participants are secured in the seat using a five-point safety harness (SCHROTH Safety Products GmbH, Arnsberg, Germany). Participants also wore a Head-and-Neck-Support (HANS) device, which consists of a helmet that is connected to a U-shaped shoulder pad via straps (SCHROTH Safety Products GmbH, Arnsberg, Germany). The helmet itself is equipped with a two-way communication system, which allows continuous contact with the experimenters as well as the playback of auditory noise during simulator motion. Fans were installed to provide additional masking of external sounds and confounding cues from wind. A photo of the setup is provided in Figure 2, and a video of (virtual) simulator motion is available as supplementary material Video S1.

**Fig. 2:**
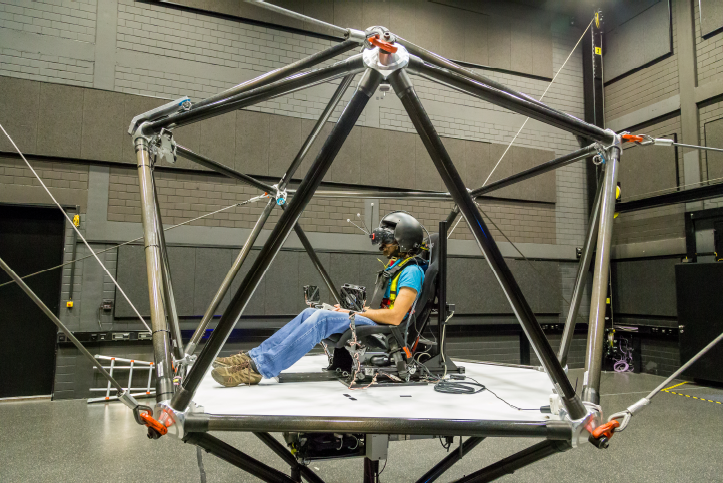
Lateral view of the experimental setup. A participant is seated in the cabin, wearing the HMD and safety equipment, and holding the button box. Fans facing the participant mask external sounds and prevent any confounding cues from wind.

Verbal responses in the Magnitude Estimation task (ME, see below) were noted by the experimenter; binary responses in the Two-Interval Forced Choice task (2IFC, see below) were registered by means of button press, using a custom made button box.

### 2.4 Task

To infer the strategy by which perceptions of traveled distance are constructed by the CNS, we would ideally measure perceptions directly and subsequently relate them to input stimuli. However, we only have access to responses on experimental tasks, which are subject to distortions that may be introduced at different stages of processing. To obtain an accurate representation of perceptions, we therefore chose to combine a Magnitude Estimation task (ME) and a Two-Interval Forced Choice task (2IFC).

In the ME task, participants were presented with visual-only, inertial-only, or combined visual-inertial motion trajectories with a range of different magnitudes (i.e., traveled distance), for which they were asked to provide verbal estimates. These verbal estimates were noted by the experimenter. This method provides an indication on the shape of the function relating perceptions to stimuli [23]. However, the method does not provide accurate information on the size of per-5 ceptual noise because direct estimates are subject to cognitive biases, such as a regression to the mean[6].

Therefore, we asked participants to also perform a 2IFC task. In the 2IFC task, participants were presented with two successive motions, designated ‘reference’ and ‘comparison’ (order randomized between trials). The reference had a constant distance, whereas the comparison had a distance that was chosen from a range of values around the reference distance. The participants’ task was to indicate which of the two was larger. Theoretically, this task is less susceptible to response biases, but it does not provide information on the shape of the distribution of the underlying perceptions.

Participants were instructed to report inertial perceptions, except when there was visual-only motion, in which case they were to report the visually traveled distance.

### 2.5 Stimuli and procedure

All motion stimuli followed a raised cosine bell velocity profile (e.g.,[20, 21, 22]). In position *X*, velocity *V*, and acceleration *A*, the profile is

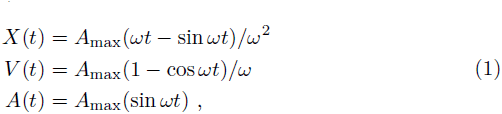

where *ω* = 2*πf*, and frequency *f* had a fixed value of 0.5Hz. Each stimulus consisted of a single period of the profile. The range of chosen *X*_max_ values was based on observations on absolute and differential thresholds in terms of *A*_max_, as reported in [25].

In the ME task, stimuli in the visual-only and inertial-only conditions had distances between 0.18m and 1.53m, evenly spaced in nine steps: [0.18, 0.35, 0.52, 0.69, 0.85, 1.02, 1.19, 1.36, 1.53]m. Each distance was presented three times, resulting in 27 visual-only, and 27 inertial-only stimuli. In the combined conditions, all possible combinations of the nine visual and nine inertial distances were presented, resulting in 81 combined trials. In total, participants performed 135 ME trials. As a familiarization with the experiment and setup, participants were presented with the range of motions from the combined condition prior to the experiment proper.

In the 2IFC task, we chose a ‘small’ and a ‘large’ reference distance, for both the visual-only and inertial-only conditions. These were 0.32m and 1.27m. The comparison distances for the small and large reference stimuli were, respectively, [0.18, 0.21, 0.25, 0.28, 0.35, 0.39, 0.42, 0.46]m, and [1.01, 1.08, 1.14, 1.21, 1.34, 1.40, 1.47, 1.53]m. We also presented participants with four combined visual-inertial conditions: these consisted of two *congruent* conditions, where visual and inertial stimuli from either the small range or the large range were paired with each other; and two *incongruent* conditions, where visual stimuli from either the small or large range were paired with inertial stimuli from the opposite range. Pairings were always done such that the n-th entry from the range of vi-6 sual distances was paired with the n-th entry from the range of inertial distances. The order of reference and comparison stimuli was also equal for visual and inertial motions, such that there was no conflict between the sensory systems with respect to correct response on the 2IFC task. To illustrate how these conditions may contribute to our understanding of perceptual mechanisms, consider the incongruent pairing of small visual and large inertial motions: if we would, for instance, observe that performance in this condition is indistinguishable from performance in the small visual-only condition, this would provide evidence for visual cue capture; alternatively, if we would observe that performance in this condition is equal to performance in the large inertial-only condition, this would provide evidence for inertial cue capture. Each reference × comparison condition was presented 10 times, resulting in a total of eight sensory conditions (two visual-only + two inertial-only + four combined) × eight comparison distances × 10 repetitions = 640 stimuli.

For an individual participant, the ME part of the experiment took approximately one hour to complete; the 2IFC part took between five and six hours to complete. The stimuli of the different sensory conditions were presented in blocks, with breaks every 15 minutes. Within these blocks, the order of stimuli was randomized. The experiment was divided into two or three sessions of two to three hours, performed on separate days. In the first session, participants first performed the ME task, followed by a part of the 2IFC task. The remainder of the 2IFC task was performed in the following session(s).

Six participants completed two additional blocks of 2IFC trials on a later date, with only congruent (small and large), multisensory stimuli, yielding 160 more stimuli. These two additional blocks were completed as a control experiment, to assess whether presenting congruent and incongruent stimuli within a single sitting a ected a-priori beliefs that visual and inertial information share a common cause. We did not find evidence for such e ects and included these data in the regular analysis.

## 3 Perception models

The models presented here are Bayesian formulations of prominent theories on how responses arise from multisensory stimulation. The models and their descriptions are adapted from [22], but include different models of sensory estimates and are expanded to more explicitly address the different stages of response generation.

We postulate that stimuli *s* yield sensory estimates *e* according to *estimate generating functions* (EGF); percepts *p* are constructed from these estimates according to *percept generating functions* (PGF); and the percepts are subsequently converted into responses *r* via *response generating functions* (RGF). Figure 3 shows a formalization of this structure. *s, e, p* and *r* are scalars that may range from *−∞*: *∞*.

**Fig. 3:**
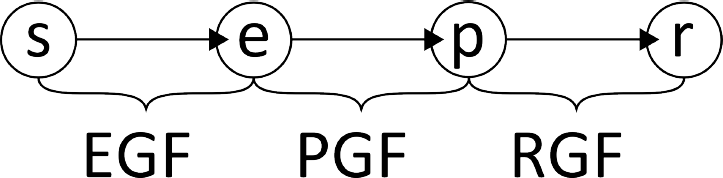
Probabilistic graphical representation of models.

The characteristics of the EGF are assumed to result from physical properties of our sensory systems, and are the same for all perception models. The PGF reflect our different hypotheses on the nature of perception, as outlined in the introduction. Finally, the RGF reflect how responses are constructed from percepts. These can incorporate knowledge of the task, and therefore differ depending on the task.

To test our hypotheses, we will fit the models directly to experimental data, and assess their fit. However, although our hypotheses mostly concern *e* and *p*, we have access only to responses *r* and stimuli *s*; *e* and *p* themselves are hidden variables. We resolve this as follows. We realize that the probability of *r* given *s* is

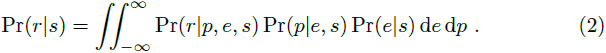

Furthermore, we assume that *p* is conditionally independent of *s* because the nervous system does not have direct access to stimuli. Similarly, we assume that the only signal available for the formation of *r* is *p*. (2) then becomes

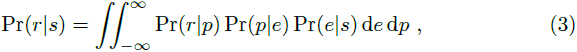

where Pr(*e|s*) is the EGF, Pr(*p|e*) is the PGF, and Pr(*r|p*) is the RGF. These functions are described separately in the following sections. We can test hypotheses on the EGF and PGF by solving the integrals, fitting the resulting models to experimental data, and subsequently comparing their fit and interpreting parameter estimates. The solution to the resulting integrals for each model of multisensory perception described in the introduction is given at the end of the Perception models section.

### 3.1 Estimate generating functions

We identify the visual (*V*) and inertial (*I*) sensory systems as sources that could provide the CNS with estimates of traveled distance. The visual system is thought to be mainly responsive to velocity of optic flow, and can generate estimates of traveled distance by integrating the vectors specified in an optic flow pattern over time. We consider the vestibular system and other sensory neurons sensitive to accelerations as a single inertial system. The CNS may obtain estimates of traveled distance on the basis of inertial information by performing a double integration.

Sensory estimates *e_m_* for modality *m* = *V*, *I*, are assumed to be noisy and possibly distorted representations of physically traveled distances *s_m_* with Gaussian noise

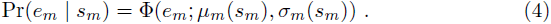

We will use Φ to represent the Normal Probability Density Function (PDF) throughout the text. Given that traveled distance represents a quantitative continuum, it is further assumed that systematic distortions that arise in the transformation of a stimulus into a sensory estimate can be described using a Stevens’ power law[23], and that the size of the noise *σ_m_*(*s_m_*) is linearly proportional to traveled distance, in accordance with a Weber’s law[26]:

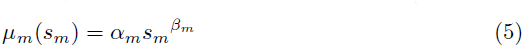

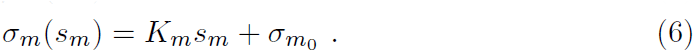

*α_m_* and *β _m_* allow the curve relating stimuli and average value of the estimates to take on a variety of shapes; 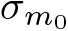 represents an absolute threshold, and *K_m_* is a Weber fraction.

### 3.2 Percept generating functions

We contend that sensory estimates of external stimuli are represented in the activity of neural populations. Neurons are tuned to respond maximally to particular stimuli, but will respond to a lesser extent to stimuli that are ‘close’. This activity over a population of neurons can then be characterized by a central estimate and a variance around this central estimate [27]. We therefore treat both *e_m_* and *σ_m_*(*s_m_*) as knowns. Furthermore, we assume that observers are unaware of the functions that relate distortions and noise to stimuli, because presence of such knowledge implies it should be compensated for. The PGF therefore use different EGF than those described in the previous section: from the observer’s perspective, the likelihood of sensory estimates given an unknown traveled distance *S* be expressed as

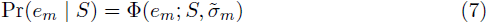

In these expressions, we use 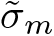 and 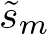 rather than *σ_m_*(*s_m_*) and *μ_m_*(*s_m_*) to emphasize that neither *s_m_* nor the relations between *s_m_* and sensory estimates are known to the CNS.

We have no particular hypotheses on how sensory estimates would be a ected by prior beliefs, and define the prior on *S* as uninformative. In the following sections, we specify how percepts *p* would be formed out of sensory estimates *e_V_*, *e_I_* according to the different models described in the introduction.

#### 3.2.1 Cue Capture models

Cue Capture (CC) models state that perception of an environmental property is dominated by a single sensory modality. In the present case, this means that multisensory perceptions are based on either the visual or the inertial estimate. Given that the prior on *S* is uninformative, the posterior probability of *S* given either sensory estimate *e_m_* is proportional to the likelihood. In accordance with the literature, we assume that the percept on an individual trial *p*_CC_ is the Maximum-A-Posteriori estimate (MAP),

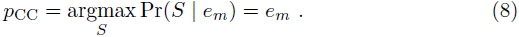

The corresponding Probability Density Function (PDF) can be expressed as

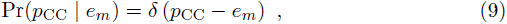

where *δ*(·) is the Dirac delta function.

We consider the above models (for the 2 *m*) special cases of a Switching Strategy (SS) model, where perception corresponds to signals provided by different modalities on a trial-to-trial basis

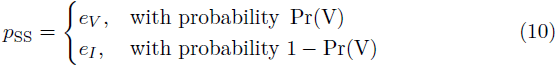

with the corresponding probability density function

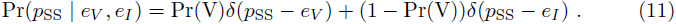

Consequently, for visual capture, Pr(V) = 1, and for inertial capture, Pr(V) = 0.

#### 3.2.2 Forced Fusion model

According to the Forced Fusion (FF) model, the CNS assumes there is a single true stimulus *S* that gave rise to both sensory estimates *e_V_*, *e_I_*. The size of this stimulus is inferred by application of Bayes’ rule. Given that the prior on *S* is uninformative, the posterior probability of *S* is proportional to the likelihood:

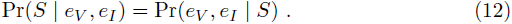

With the additional assumption of independence of the estimates, this becomes

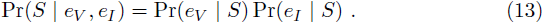

As a percept *p*_FF_, we again choose the MAP estimate:

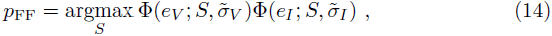

which yields the expression (see e.g., [12])

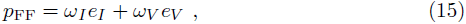

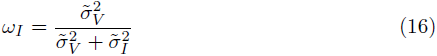

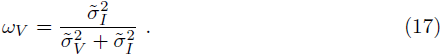

Similar to (9), the PDF can be written as

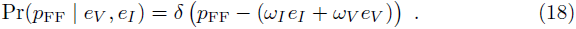

#### 3.2.3 Causal Inference model

Causal Inference models differ from cue combination models in that it is not assumed that multisensory information always shares a common cause. Instead, inferences are made on the causality of multisensory information and these a ect subsequent processing of the sensory estimates in the formation of percepts.

In the present implementation, the percept *p*_CI_ is a weighted average of *p*_FF_ and *p*_SS_, with weights proportional to the probability that the sensory estimates share a common cause Pr(C | *e_V_*, *e_I_*), and its complement that they do not 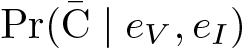:

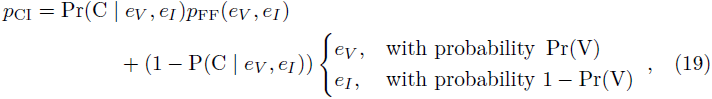

This formulation reflects the Model Averaging CI strategy (e.g., [18, 28, 29]). It is similar to earlier work by our group on heading estimation [20, 30, 21] and the perception of verticality [22].

The probability of a common cause given the sensory estimates is

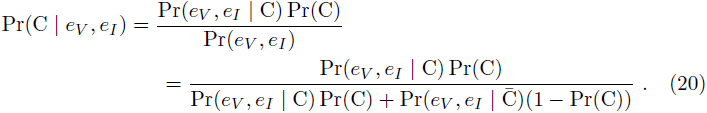

Pr(C) is a free parameter that represents a participant’s a-priori belief that signals will share a common cause. The likelihood of the sensory estimates *e_V_*, *e_I_* given a common cause C is

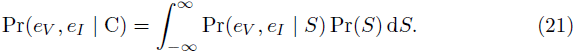

With the prior on *S* again assumed uninformative, this becomes

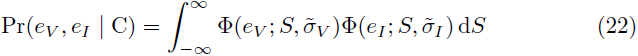

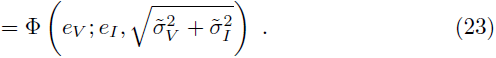

For an interpretation of independent causes, a percept will be based on either the visual or the inertial estimate. Assuming independence, this means that 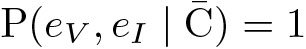.

Consequently,

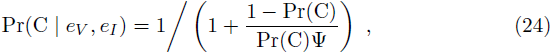

with Ψ as a placeholder for equation (23).

The PDF of *p*_CI_ given *e_V_*, *e_I_* can be expressed as

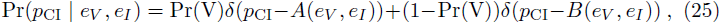

where

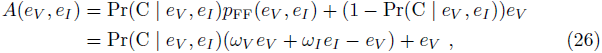

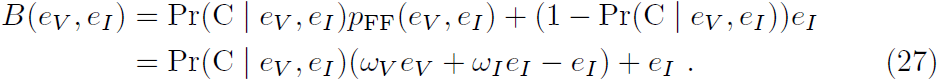

### 3.3 Response generating functions

The RGF specify how responses relate to perceptions. Because the types of and constraints on responses depend on the task, the RGF are specified separately for the ME and 2IFC tasks.

#### 3.3.1 Magnitude Estimation task

For the ME task, we assume that the actual response *r*_ME_ is based upon the percept *p*_model_, but is subjected to additional distortions that reflect participants’ a-priori beliefs on the probability of certain responses being a correct representation of the stimulus.

For the present case, we expect that participants may be inclined to report the mean magnitude learned during the training phase of the experiment *µ*_0_^1^, such that

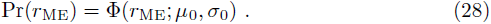

The likelihood of a percept given a response is modeled as

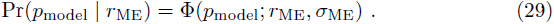

This yields the posterior

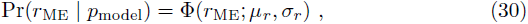

with

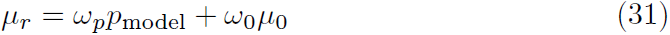

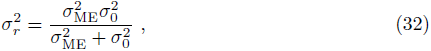

where

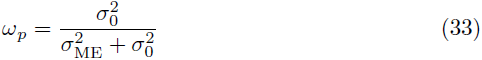

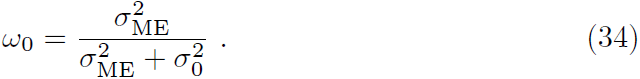

#### 3.3.2 Two-Interval Forced Choice task

On individual trials, participants respond positively(*r*_2IFC_ = 1) if the perceived distance for the comparison stimulus *p_C_* exceeds the perceived distance for the reference stimulus *p_F_*; and they give the opposite response (*r*_2IFC_ = 0) otherwise.

We assume that this binary response is not a ected by prior beliefs.

The probability of a positive response is equal to the area under the surface specified by the joint density of *p_F_*, *p_C_*, for which *p_C_* > *p_F_* (i.e., above the diagonal).

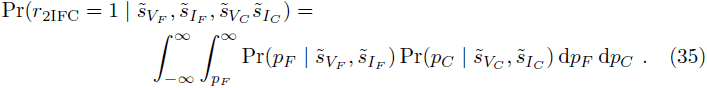

The probability of the alternative response Pr(*r*_2IFC_ = 0 | *p_F_*, *p_C_*) is the complement of equation (35). These integrals were calculated numerically for each model.

### 3.4 Model expressions

Using the expressions for the EGF, PGF and RGF, we can derive the expressions Pr(*r*_model_ | *s*) and Pr(*p*_model_ | *s*) needed to fit the models to the data of the ME and 2IFC tasks, respectively.

For the CC*_m_* (with *m* = *V*, *I*) models, we obtain:

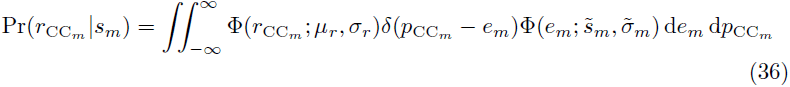

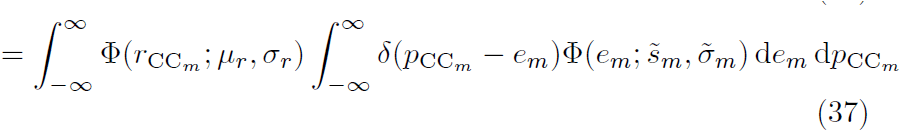

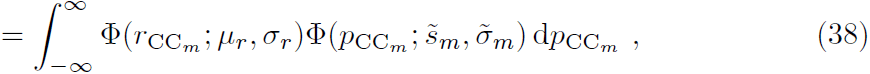

with *m* referring to the particular modality chosen.

The latter probability density, 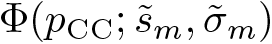, is the expression used to calculate the probability of reference and comparison percepts on the 2IFC task, needed to calculate the probability of a positive response.

Solving the integral in (38) yields the expression for the ME task:

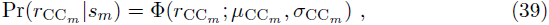

where

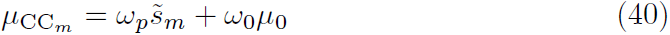

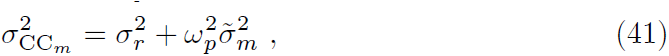

and 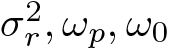 as in (32), (33), and (34).

By combining the two possible CC models, we obtain the expressions for the SS model

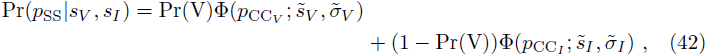

and

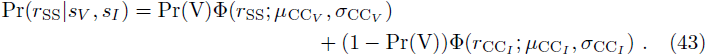

By following the same steps, we obtain the expressions for the FF model

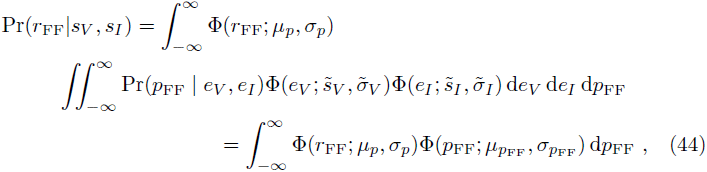

with

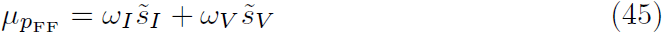

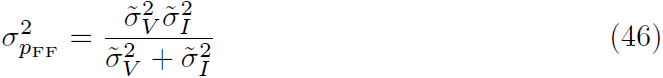

with *ω_I_*, *ω_V_* as in (16) and (17). As in (38), the second PDF inside the integral in (44) is the expression for the probability of perceptions *p_F_*, *p_C_*, used to calculate the probability of a positive response on the 2IFC task. Solving the integral yields

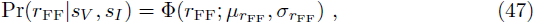

where

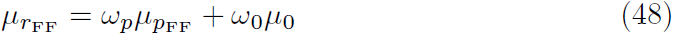

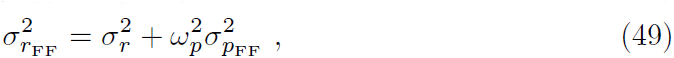

Put simply, according to the CC and FF models, responses on the ME task are a weighted sum of percepts and a belief about possible responses *µ*_0_.

Finally, for the CI model, we obtain:

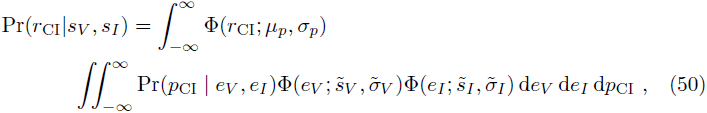

with Pr(*p*_CI_ | *e_V_*, *e_I_*) as in (25):

Due to the expression for Pr(C | *e_V_*, *e_I_*) in *A*(*e_V_*, *e_I_*) and *B*(*e_V_*, *e_I_*), there is no closed-form solution to (50). This can be dealt with by linearizing *A*(*e_V_*, *e_I_*) and 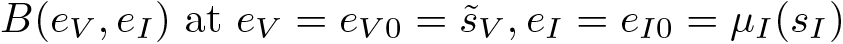, as described in [22]:

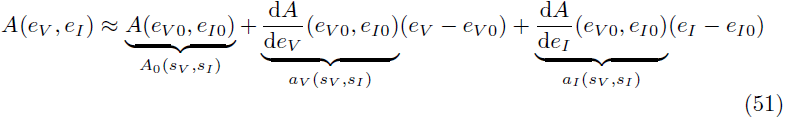

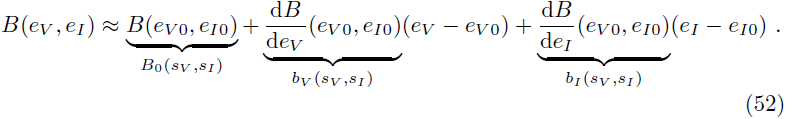

Using the approximation, the integrals can be solved. This yields the following expression for the 2IFC task:

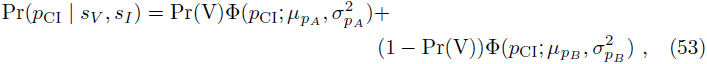

with

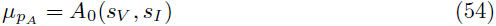

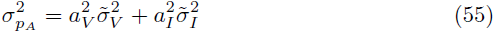

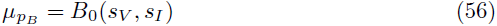

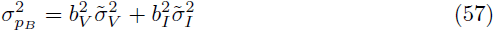

and for the ME task:

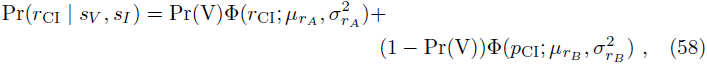

with

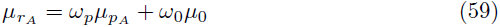

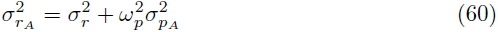

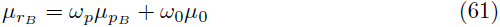

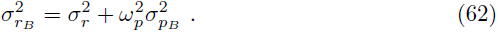

*ω_p_*, *ω*_0_ in these latter equations are as in (33) and (34).

### 3.5 Model fitting and Data analysis

The parameters to account for systematic distortions in sensory estimates (*α_V_*, *β_V_*, *α_I_*, *β_I_*), the size of the noises 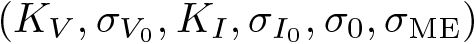, and the dominance of visual information and common cause priors (Pr(V), Pr(C)) for multisensory perception were treated as free parameters, yielding a total of 10-12 free parameters, depending on the model.

It should be noted that a comprehensive model of perception would ideally account for unisensory and multisensory conditions in the same way. However, we deviated from this notion because there were differences in the task for the different conditions: in visual-only conditions, participants were instructed to report traveled distance as suggested by the visual stimulus, whereas they were instructed to report physical distance in the other conditions. In the inertial-only conditions the HMD showed black screens, which provides no information, rather than a stationary scene which indicates standstill. Therefore, the expressions for visual- and inertial capture CC*_V_*, CC*_I_* were used to model responses in the corresponding unisensory conditions for all models.

The models were fitted by minimizing the combined negative log-likelihood nLL of the responses to the two tasks given the models:

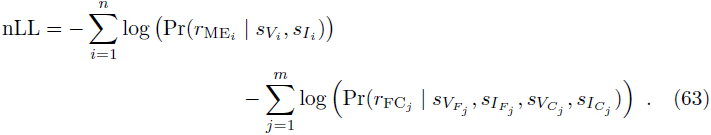

optimizations were performed using the fmincon routine in MATLAB. As sanity checks, we 1. made sure that the CI model yielded identical nLL scores as each alternative model when it was passed the parameters found for that model, and 2. we made sure that the nLL score of the SS model was always lower than either CC model, and that the nLL score of the CI model was always lower than either other model.

Conclusions on which model best explained data were made on the basis of the Bayesian Information Criterion (BIC) scores [31], which are penalized likelihood scores. Smaller BIC scores indicate better performance. A difference between BIC scores (ΔBIC) of 0-2 is considered negligible evidence; 2-6 is positive evidence; 6-10 is strong evidence; and > 10 is considered decisive evidence.

## 4 Results

### 4.1 Model comparisons

To test our hypotheses, we presented participants with both unisensory visual-only and inertial-only motions, as well as with multisensory visual-inertial motions with a range of discrepancies. Participants performed two tasks: in the first part of the experiment, they provided verbal estimates of traveled distance (ME). In the second part of the experiment, participants provided binary responses on which motion had the larger displacement for numerous pairs of motions with various discrepancies (2IFC). We assume that both tasks rely on the same percepts, and we fitted the models described in the Methods and Materials section to the data of both tasks simultaneously. As an illustration of the data and model fits, Figures 4 and 5 below respectively show data from the ME and 2IFC task and the CI model fit for an example participant.

**Fig. 4:**
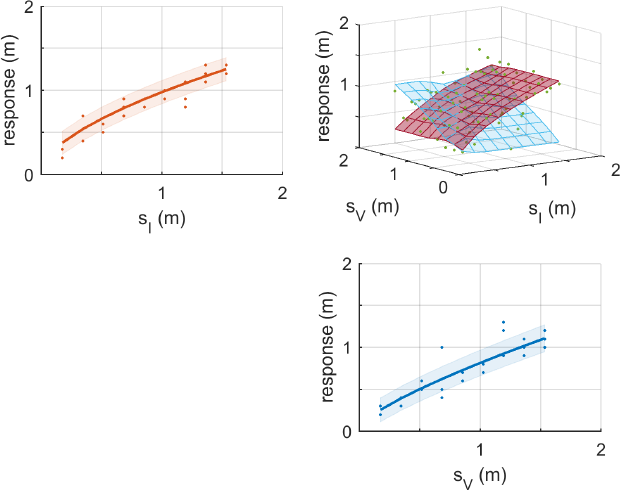
Data and CI model fits from the ME task for an example participant (9). Dots represent individual responses; lines and surfaces represent the mean responses as obtained from the CI-model fit; shaded areas around the lines show 1 standard deviation 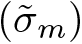. Upper left/lower right panel: responses in the inertial-only (orange) resp. visual-only (blue) conditions plotted against stimulus distance *s_V_* resp. *s_I_*. Upper right panel: responses in the combined condition plotted against stimulus distances *s_V_, s_I_*. The light blue and red surfaces respectively correspond to the branch of the CI model where percepts are constructed as a weighted average of *p*FF and *p*CC*_V_*, or *p*FF and *p*CC*_I_*. Note how the surfaces of the branches appear to converge for small discrepancies, reflecting FF; and to diverge for large discrepancies (CC).

**Fig. 5:**
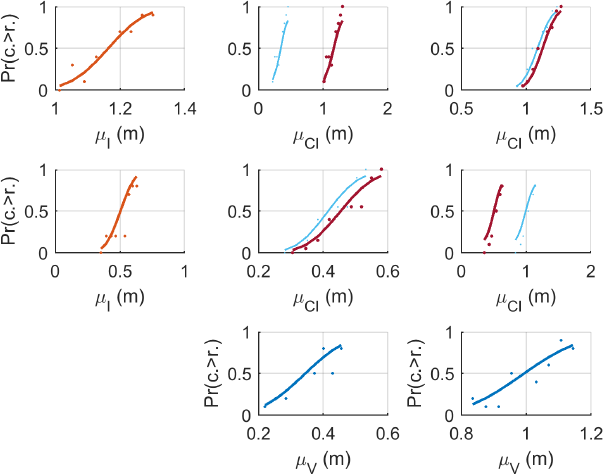
Data and CI model fits from the 2IFC task for an example participant (9). Dots represent the proportion of positive responses for a given reference × comparison condition; lines represent the probability of a positive response as obtained from the CI-model fit, plotted as functions of the *perceived* distances obtained from the joint fit of the model to the data of the ME and 2IFC tasks. The left column of panels shows the data and fits for the large (top row) and small (middle row) inertial-only (orange) conditions; the bottom row of panels shows this for the small (middle column) and large (right column) visual-only (blue) conditions. The other panels show the multisensory conditions specified by the corresponding column and row. For these conditions, the light blue line represents the CI model branch where percepts are constructed as a weighted average of *p*FF and *p*CC*_V_*; the red lines correspond to the branch combining *p*FF and *p*CC*_I_*. The size of the circles in the multisensory plots is proportional to parameter Pr(V), indicating that inertial information was dominant for this participant. Note how the psychometric functions for the two branches of the model are similar in the congruent multisensory condition, whereas they diverge in the incongruent conditions.

In Table 1, we present the obtained ΔBIC scores for the individual fits as well as overall ΔBIC scores for each model. Individuals show considerable variability with respect to which model best explains their data. A variant of a cue capture model (CC*_m_* or SS) provides the best explanation for 6/9 participants; the FF model provided the best explanation for 2/9 participants, and the CI model provides the best explanation for 1 participant. However, taken together, the CI model best explains the data, with evidence that may be considered decisive compared to the other models: ΔBIC = 55.37, relative to runner-up model SS. A complete overview of all model fit indices is provided in supplementary material Table S1.

**Table 1:**
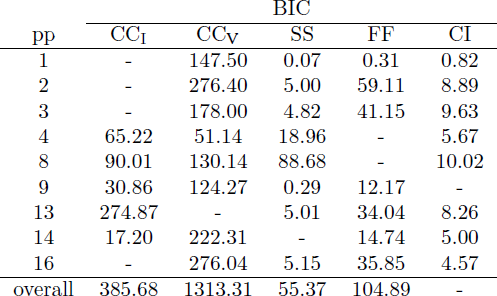
ΔBIC scores for all models (i.e., BIC_*model*_ − BIC*_best_*), for each participant (pp). For each model, overall BIC scores were calculated using the sums (over participants) of the model log-likelihoods, the number of parameters, and the total number of observations.

| pp | BIC |  |  |  |  |
| --- | --- | --- | --- | --- | --- |
| | $\text{CC}_I$ | $\text{CC}_V$ | SS | FF | CI |
| 1 | - | 147.50 | 0.07 | 0.31 | 0.82 |
| 2 | - | 276.40 | 5.00 | 59.11 | 8.89 |
| 3 | - | 178.00 | 4.82 | 41.15 | 9.63 |
| 4 | 65.22 | 51.14 | 18.96 | - | 5.67 |
| 8 | 90.01 | 130.14 | 88.68 | - | 10.02 |
| 9 | 30.86 | 124.27 | 0.29 | 12.17 | - |
| 13 | 274.87 | - | 5.01 | 34.04 | 8.26 |
| 14 | 17.20 | 222.31 | - | 14.74 | 5.00 |
| 16 | - | 276.04 | 5.15 | 35.85 | 4.57 |
| overall | 385.68 | 1313.31 | 55.37 | 104.89 | - |

### 4.2 Parameter estimates

Estimates of model parameters (Table 2) were generally consistent among models, insofar as the parameters were common to models. In the following, we therefore consider the parameter estimates of the CI model, as it contains all parameters found in the models.

**Table 2:** Parameter estimates for the CI model for each individual and the median value across participants. Estimates of parameter values were highly consistent among models. See the main text for a description of the parameters.

| pp | parameter | | | | | | | | | | $\text{Pr}(V)$ | $\text{Pr}(C)$ |
| --- | --- | --- | --- | --- | --- | --- | --- | --- | --- | --- | --- | --- |
| | $\alpha_I$ | $\beta_I$ | $K_I$ | $\sigma_{0_I}$ | $\alpha_V$ | $\beta_V$ | $K_V$ | $\sigma_{0_V}$ | $\sigma_0$ | $\sigma_{ME}$ | | |
| 1 | 1.48 | 0.67 | 0.00 | 0.13 | 0.99 | 1.28 | 0.16 | 0.06 | 0.57 | 0.23 | 0.02 | 0.97 |
| 2 | 0.91 | 1.12 | 0.11 | 0.05 | 0.93 | 1.53 | 0.19 | 0.00 | 0.12 | 0.03 | 0.00 | 0.13 |
| 3 | 1.58 | 0.82 | 0.01 | 0.18 | 0.93 | 1.40 | 0.20 | 0.00 | 0.36 | 0.21 | 0.00 | 0.00 |
| 4 | 0.75 | 1.11 | 0.32 | 0.01 | 0.61 | 1.35 | 0.05 | 0.12 | 0.22 | 0.10 | 0.00 | 0.98 |
| 8 | 1.01 | 0.62 | 0.00 | 0.14 | 0.98 | 1.30 | 0.04 | 0.15 | 0.24 | 0.10 | 0.50 | 1.00 |
| 9 | 1.00 | 0.61 | 0.00 | 0.06 | 0.83 | 0.76 | 0.03 | 0.07 | 0.44 | 0.13 | 0.13 | 0.73 |
| 13 | 0.50 | 1.33 | 0.14 | 0.00 | 0.52 | 1.63 | 0.13 | 0.00 | 0.19 | 0.10 | 1.00 | 0.15 |
| 14 | 0.97 | 1.14 | 0.05 | 0.05 | 0.75 | 1.42 | 0.02 | 0.11 | 0.33 | 0.11 | 0.09 | 0.00 |
| 16 | 0.74 | 1.19 | 0.14 | 0.02 | 0.62 | 1.47 | 0.10 | 0.03 | 0.00 | 0.00 | 0.00 | 0.33 |
| median | 0.98 | 1.11 | 0.05 | 0.05 | 0.83 | 1.40 | 0.10 | 0.06 | 0.24 | 0.10 | 0.02 | 0.33 |

Median values of parameters that describe inertial distortions were *α*_*I*_ = 0.98, *β*_*I*_ = 1.11, indicating that inertial distances were somewhat underestimated, although the underestimation decreases with stimulus distance and eventually reverses, becoming overestimation after about 1.2m. Parameters that describe inertial noise as a function of stimulus distance *K_I_* = 0.05 and 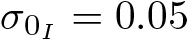 indicate that noise increases with distance.

Visually, distances are also initially underestimated and eventually overestimated, with a reversal around 1.5m (*α_V_* = 0.83, *β_V_* = 1.40). Visual noise also increased with stimulus distance, with median parameter values *K_V_* = 0.10, 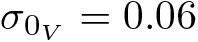.

Parameter *σ*_0_ is the standard deviation of the prior belief on percepts reflecting the median stimulus, which comes to expression as the strength of the regression to the mean e ect in the ME data. The median value of this parameter was 0.24. This value is several times larger than the standard deviations of the sensory estimates.

*σ*_ME_ reflects the standard deviation of the ME task, which is additional noise that is introduced in the formation of responses rather than at the level of sensory estimation. The median value of this parameter was 0.10.

Parameter Pr(V) reflects the degree of visual dominance. The median value was 0.02, showing that participants were mostly able to focus on the inertial motion, as per instructions. Debriefing after the experiment indicated that the case where the parameter was close to 1 was caused by misinterpretation of the instructions.

Parameter Pr(C), which reflects an a-priori belief that signals will share a common cause (regardless of any sensory information), had a median value of 0.33, indicating that participants were a-priori more likely to decide that multisensory stimuli had separate rather than common causes.

A complete overview of all parameter estimates for all models is provided in supplementary material Table S2.

## 5 Discussion

The main question which the present study aimed to address was whether processing of quantitative multisensory information on self-motion can be adequately described using a CI model. CI differs from models that assume that multisensory percepts are constructed as a weighted average (FF) or by dominance of a single modality (CC, SS) by explicitly including an assessment of causality of sensory estimates in the formation of percepts. Put simply, we investigated if the CNS determines whether or not multisensory information should be integrated.

In addition, we combined tasks designed to either obtain information on accuracy and precision of perception. This was done to disentangle distortions introduced at different levels of processing. In the following sections, we discuss our findings on distortions in perception and on the applicability of the CI model, and relate the findings to the literature.

### 5.1 Unisensory distance estimation

Traditionally, distortions in the accuracy of responses on estimation tasks have been attributed to perception following logarithmic [26] or power laws [23]. In contrast, in more recent models of perception based on Bayesian inference[32, 12, 33, 13], distortions are, generally, attributed to interactions between (unbiased) sensory estimates and prior knowledge. Although these approaches can be unified [6], it begs the question whether systematic distortions in perception are caused by our sensory organs, by interaction between sensory estimates and prior knowledge, or both. We aimed to address this question by using a combination of an ME task and a 2IFC task. The ME task provides information on the accuracy of percepts by allowing us to characterize the shape of the response function, and it provides information on the precision of perception by revealing the response distribution. However, it may introduce additional noise and distortions caused by any cognitive strategies participants may apply. In contrast, the 2IFC task should not be subject to such additional noise and distortions, but the task only provides information on the precision of perception; not on the shape of the underlying distribution of percepts. By combining the ME and 2IFC tasks and formulating models such that the same percepts underlie responses on both tasks while accounting for any additional distortions in the ME task, we aimed to quantify both sensory distortions and e ects of prior knowledge.

We used Stevens’ and Weber’s laws as templates for accuracy and precision of sensory estimates, respectively. The data gathered in the unisensory conditions allowed us to characterize systematic distortions of sensory estimates compared to the stimuli, and to determine absolute detection and differential thresholds for visual and inertial distance estimation. These characterizations then provided benchmarks for comparison of the present data with prior research on visual and inertial perceptions of self-motion[34, 35, 25, 36, 37].

For inertial distance estimation, the Stevens’ power law parameter estimates had median values of *α_I_* = 0.97 (range 0.50-1.58), and *β_I_* = 1.11 (range 0.61-1.19). The *α* parameter can be interpreted as a gain and the *β* parameter determines whether the scaling is constant or varies with traveled distance. The observed range of values indicates that there is considerable variability between individuals in how inertial motion is perceived. Nevertheless, the median values of the parameters indicate that sensory estimates of traveled distance show underestimation, but that this underestimation eventually changes into overestimation at 1.2m.

The findings on inertial sensory noise indicate a median absolute detection threshold 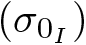 of 0.05m, which corresponds to velocity and acceleration detection thresholds of 0.05m/s and 0.08m/s^2^ (for the present motion profile). Thresholds for motions along the antero-posterior axis reported in the literature are in the same order of magnitude, but generally somewhat smaller: Benson [34] notes a 67% correct-detection threshold of 0.06m/s^2^ for a 0.3Hz cosine-bell profile; Zaickik et al., [35] find a value of approximately 0.03m/s^2^ for detection of the direction of sinusoidal fore-aft motion.

Naseri & Grant [25] determined 70.7% differential thresholds for 0.4Hz and 0.6Hz motions, with maximum accelerations of 0.5, 1.0, 1.5, and 2.0m/s^2^. By interpolation, we find 0.5Hz thresholds that amount to 0.060, 0.077, 0.094, and 0.111m/s^2^. We can approximate 70.7% differential thresholds in terms of acceleration from the present data. This yields values of 0.059, 0.071, 0.083, and 0.096m/s^2^, which are strikingly similar to the values reported be Naseri & Grant^2^.

For estimates of traveled distance based upon visual information, we obtained median values of *α_V_* = 0.83 (range 0.52-0.99), and *β_V_* = 1.40 (range 0.76-1.63). Again, we note considerable interpersonal variability. The median parameter estimates suggests that visual estimates of distance are also underestimated, but that this underestimation decreases with visually traveled distance, changing from under- to overestimation at 1.5m.

Literature on visually induced perception of linear motion [36, 37] (i.e., linear vection) indicates a lower boundary on optic flow velocity of 0.03m/s for the sensation to be induced, which can be interpreted as an absolute detection threshold. Using the observed median value for 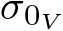 (0.06), we find a similar velocity threshold of 0.06m/s for the present experiment. Literature on differential thresholds for vection is scarce. However, Sauvan & Bonnet [38] note a Weber fraction of 41% for 30s constant-velocity motion profiles. This value is considerably higher than in the present study (*K_V_* = 0.10(10%)). It is likely that the discrepancy is due to the fact that the present stimuli had very different motion profiles, and were not specifically designed to assess vection.

Parameter *σ*_ME_ expresses additional noise in responses introduced by the ME task, as compared to the 2IFC task. The median value of this parameter was 0.10, indicating that the additional noise in the ME task is comparable in magnitude to the sensory noise. However, we again note considerable interpersonal variability (range 0.00-0.23), which indicates that some participants were able to express the perceived distance very precisely, whereas others were not. It is possible that these values reflect individual differences in round-off tendencies. Finally, the observed median value for model parameter *σ*_0_ was 0.24 (range 0.00-0.57). In comparison to *σ*_ME_, this parameter can be interpreted as expressing the strength of the regression to the mean e ect[6]. Filling in the median values in equations (33) and (34), we find that the relative contributions of the perceived distance and the prior belief that the traveled distance is the mean traveled distance are 0.85: 0.15. This indicates that whereas the regression to the mean e ect is small, bias in responses is introduced both at the sensory level and by high-level processing.

### 5.2 Multisensory distance estimation

Overall, the model comparisons provide decisive evidence for the CI model over the alternatives. However, on an individual level, the results of the analysis are more varied; favoring different models as the best explanation of data of individual participants, and predominantly a variant of the CC/SS models. This variability could be interpreted as a sign that individuals are wired differently with respect to processes governing the formation of perceptions. However, we believe this to be unlikely: a similar variability has been reported in recent studies on the applicability of CI for perception of heading [20, 21] and verticality [22]. As in the present study, it was found that different models were preferred over CI in individual cases, although in these studies the FF model was preferred in the majority of cases. By increasing the range of discrepancies, segregation behavior became more apparent, and it was established that participants were indeed likely to perform causal inference. In contrast, segregation behavior was already apparent in the present study. In fact, it is arguably unambiguous because all participants were able to report visual distances in the visual-only conditions, and physical distances in the inertial-only conditions. This also implies that in any study on self-motion perception where unisensory conditions are employed, it is implicitly assumed that participants are able to segregate multisensory information. An interpretation of the results in favor of a variant of a cue capture model would also be inconsistent with those cases where FF was observed, as well as with the findings of studies showing integration [14, 15]. The CI model accounts for interpersonal differences by assuming that the tolerance for discrepancies is an idiosyncratic property. This is assumption is essentially not different from assuming that accuracy and precision of sensory estimates are idiosyncratic properties.

A second argument to further consider the CI model comes from vehicle simulation research. Studies in this field have established so-called Coherence Zones (CZ). A CZ is defined as a fixed range of visual-inertial motions for which participants accept the common cause hypothesis, despite some physical discrepancy. The concept was derived from the observation that physically moving observers are unable to detect concurrent visual motion when the amplitude of the visual motion is below a particular threshold [39, 40]. CZ have been determined for the amplitude and phase of rotational as well as translation motion [41, 42, 43, 44, 45], and also for heading [46, 30]. CI models can be said to be formed around an analogous concept, namely the probability that visual and inertial motion with any discrepancy will be attributed to a common cause Pr(C | *e_V_*, *e_I_*). Hence, despite CZ not being concerned with mechanisms governing perception per se, they do provide benchmarks for studies on CI.

As an illustration, Figure 6 shows in the left panel probability Pr(C | *e_V_*, *e_I_*) as obtained from the CI model, and in the right panel the CZ for surge-motion from the literature [41]. A formal comparison between the two concepts is problematic, because [41] do not specify the targeted percentage-correct threshold for the determination of the CZ. Regardless, the resemblance between the CZ and the probability map obtained from CI modeling is evident. Moreover, according to the CI model, the peakedness of Pr(C | *e_V_*, *e_I_*) results from the sum of the visual and inertial variances (in combination with the a-priori belief in a common cause; see section 3.2.3). In other words, the width of the band for which Pr(C | *e_V_*, *e_I_*) is relatively high is predicted by the sum of the sensory variances. This theoretical result corresponds to empirical findings on the width of the CZ for yaw rotations[45].

**Fig. 6:**
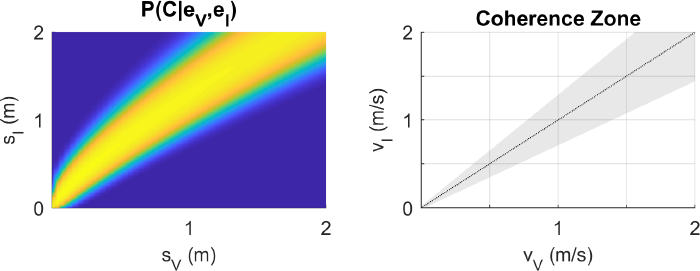
Left panel: probability of a common cause judgment given perceptions *e_V_, e_I_* (Pr(C = 1|*e_V_, e_I_*)), for stimuli with visual and inertial traveled distances (*e_V_, e_I_*) between 0 *−* 2m for an example participant (9). Blue indicates low probability; yellow indicates high probability. The curvature of the band of high probability is due to the inclusion of distortions in the modeling of sensory estimates. Right panel: coherence zone (CZ) plot, based on the function for forward surge motion specified in [41].

In conclusion, we have demonstrated how different causes of sensory bias may be disentangled and quantified using a combination of ME and 2IFC tasks. Although definitive evidence for CI was not obtained, we have also shown that perception of distance of horizontal linear motion is consistent the notion that the CNS includes assessments of signal causality in the formation of perceptions, and that this notion accounts for empirical findings in related fields.

## 6 Acknowledgments

The authors thank Dr.med. Ikar Lau er for medical supervision of the experiment and Mikhail Katliar for reviewing the mathematics. The virtual environment used in the present study was developed by Dr. Martin Breidt and Daniel Diers for the “Origins – steps of humankind” exhibition, running from May 20, 2017 to December 3, 2017, in Schlossmuseum Tübingen, Germany.

## Footnotes

1 As a mean for the prior, we also tested a value of 0, which reflects a prior tendency to report ‘no motion’. The fits were however less good, as evidenced by higher BIC values.

2 These thresholds were determined as follows: first, *µ_I_* (*x_I_*) and *σ_I_* (*x_I_*) were calculated according to the functions described in section 3.1, using the median parameter estimates and stimulus accelerations from [25] converted to meters. The obtained values are used as the mean and standard deviation parameter of cumulative Gaussians as approximations of the perception model output for the 2IFC task. We then calculate the 70.7% thresholds using the inverse of this function, subtract *µ_I_*, and convert the resulting values back to accelerations.

